# Prediction of Direct Carbon Emissions of Chinese Provincial Residents under Artificial Neural Networks in Deep Learning Environment

**DOI:** 10.1101/2020.07.14.202119

**Authors:** Jin Hui

## Abstract

It is aimed to deepen the understanding of the consumption carbon emissions of Chinese provinces, establish an accurate and feasible carbon emission prediction model, develop an urban low-carbon economy, and ensure the sustainable development of Chinese cities. Through the national statistical data information, based on the artificial neural network model, mathematical statistics and deep learning methods are used to learn and analyze the carbon emission data of various provinces in China from 1999 to 2019. The neural network toolbox in Matlab is used to program separately to realize the prediction of carbon emissions by different neural network models. After comparing and analyzing the accuracy and prediction performance, the optimal model for the prediction effect is selected. Finally, based on ArcGIS Engine (Arc Geographic Information Science Engine) and C#.NET platform, the call to Matlab neural network toolbox is realized. The selected model is embedded in the prediction system to complete the development of the entire system. The results show that the carbon emissions of residents in the north are distinctly higher than those in the south. Also, with the passage of time, the rate of carbon emissions continues to accelerate. Compared with other models, Elman neural network has higher accuracy and smaller error in carbon emission prediction. Compared to BP (Back Propagation) neural network, the accuracy is improved by 55.93%, and the prediction performance is improved by 19.48%. The prediction results show that China is expected to reach the peak of carbon emissions from 2027 to 2032. This investigation will provide a theoretical basis to control and plan carbon emissions from Chinese urban residents.

## 1. Introduction

Recently, with the continuous expansion of the field of human activities, the ecology of nature is facing many problems. A series of environmental problems such as global warming, frequent extreme weather, and melting glaciers threaten human survival and health [1]. Among them, global warming has become an important issue of common concern in today’s society and related investigations have also emerged endlessly [2]. It is found that the main factors affecting climate warming come from the continuous emission of CO_2_ [3]. To reduce carbon emissions, countries have implemented the most stringent measures [4]. China is the largest developing country in the world today and one of the major energy-consuming countries [5]. As a big country that takes charge of the human living environment, China has the responsibility to actively undertake emission reduction tasks according to its ability while doing well in national economic development [6]. The carbon emissions brought about by rapid urbanization and industrialization as well as the direct carbon consumption of residential energy have become the main part of China’s greenhouse gas emissions [7]. Relevant investigations show that in developed countries, as the industrial level continues to decrease, more residents in cities continue to increase their carbon emissions. Some regions have exceeded industrial carbon emissions [8]. Therefore, according to the current energy consumption and carbon consumption levels of residents in various provinces of China, it is greatly significant to predict the direct carbon emissions of Chinese residents.

The neural network is a newly developed computer technology in recent years. It relies on its unique network structure characteristics and data processing methods to achieve fruitful results in many fields [9]. It includes engineering automation, image recognition, model prediction, and signal processing. Among them, the use of neural networks to build prediction models is a relatively common method [10]. There have been many investigations on the application of neural network prediction models in carbon emissions. Among them, Ye et al. (2018) used neural networks to predict carbon emissions in the construction industry. It was found that the economic development of the construction industry and the improvement of standards may have a significant impact on future carbon dioxide emissions [11]. Balki et al. (2018) used back-propagation artificial neural networks to develop models that can estimate engine performance and exhaust emissions. It was found that the carbon emissions of ethanol vehicles were reduced by 6% compared to gasoline vehicles [12]. Liu et al. (2017) used chaos theory combined with BP (Back Propagation) neural network to fit and predict carbon emission time series without considering other factors. It was easier and more accurate than other prediction methods [13]. Kim et al. (2017) used the newly developed neural network model to predict and store with high accuracy. It can effectively evaluate the carbon emissions from industry and residents in the deep brine [14]. From the above works, it can be seen that the use of neural network models in carbon emissions prediction has been one of the hot issues in this field. But the relevant works almost revolve around automobiles, buildings, and industries in developed countries. Few works have involved predictions of residential carbon emissions.

The intelligent methods are innovatively used to explore residents’ direct carbon emissions in the investigation. Different neural network models are used to conduct a comparative investigation in terms of the error between the true value and the predicted value, the residual error of the prediction result, and the MSE (Mean Squared Error). Through the use of mathematical statistics and deep learning methods, the carbon emission data of each province in 1999-2019 are learned and analyzed. Based on a common language development platform, GIS (Geographic Information Science) components are combined and carbon emission prediction models are embedded to design a comprehensive and integrated visual intelligent platform. The platform can be developed to predict the direct carbon emissions of Chinese residents. It provides a theoretical basis for investigations related to China’s carbon emissions and provides a scientific basis for the control and planning of residents’ carbon emissions.

## 2. Methods

### 2.1 Calculation method of carbon emissions

Carbon emissions can be divided into direct carbon emission calculation and indirect carbon emission calculation. The direct carbon emission represents the carbon emission generated by residents in the process of direct energy consumption. It mainly comes from the energy consumption of residents, including carbon emissions from coal, natural gas, electricity, oil, and geothermal heat [15]. The calculation method is mainly to use the carbon emission coefficient method, that is, to calculate the carbon emissions of residents’ lives based on the statistical data of various fossil energy sources. This investigation refers to the calculation method of international greenhouse gas emissions [16], and the calculation equation is as follows.

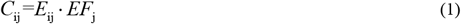

Where: i is the i-th region and j is the j-th fuel. When j is 1, 2, 3…19, it represents 19 kinds of energy sources such as coal, oil, natural gas, heat, and electricity. *C*_ij_ is the carbon emission of the j-th fuel in the i-th region. *E*_ij_ is the final consumption of the j-th fuel in the i-th region. *EF*_j_ is the CO_2_ emission coefficient of the j-th fuel. This method needs to count plenty of residents’ energy consumption data, which takes a long time.

Indirect carbon emission refers to judging the consumption of energy by residents’ consumption in their lives and calculating carbon emissions. It mainly corresponds to residents’ indirect energy consumption. Commonly used models mainly include the input-output model, life cycle assessment method, hybrid life cycle method, and consumer lifestyle method [17]. The calculation equation of the input-output model is as follows.

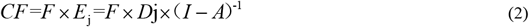

Where: CF represents indirect carbon emissions. F represents the living consumption of residents. Ej represents the indirect carbon emission intensity of energy consumption in sector j. D_j_ represents the direct carbon emission intensity of energy consumption in sector j. A represents the matrix of input and output direct consumption coefficients. I represents the identity matrix of the same order of A. However, the scope of the model is wider and there are more sectors. Therefore, the period of preparation and calculation is longer, and there are greater limitations.

The life cycle assessment method mainly evaluates the overall processing, manufacturing, transportation, and sales of a certain commodity to calculate carbon emissions. Compared with the input-output model, it needs to obtain plenty of business data. Therefore, the model is more difficult to implement. The hybrid life cycle method is a combination of input-output and life cycle, considering the micro-scale adaptability of input-output, and reducing the dependence on business data. The main calculation equation is as follows.

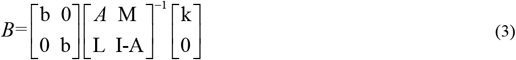

Where: B represents the carbon emission of a product. b represents the matrix of microscopic carbon emission coefficients. A represents the technical matrix. I represents the identity matrix. L represents the input of the macroeconomic system to the microsystem of the commodity. M represents the input of the commodity microsystem to the macroeconomic system. K represents the external demand.

The consumer lifestyle method is a method that uses lifestyle to explore the relationship between consumer activities and the environment. It can calculate carbon emissions by measuring various consumption data. The calculation equation is as follows.

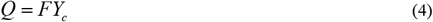

Where: F represents a row vector, which means the intensity of carbon emissions. Y_c_ represents a column vector, which means the household consumption expenditure. This investigation focuses on the development trend and prediction methods of carbon emissions. Also, indirect methods have various drawbacks. Therefore, in the calculation of carbon emissions, the method of direct carbon emissions is selected.

### 2.2 Data source and indicator measurement of carbon emission prediction

The residents’ energy consumption structure is diversified. Different regions and households have different types of energy consumption. For the calculation of carbon emissions, most provinces and cities are used as administrative units [18]. In this investigation, the carbon emission coefficient method is used to calculate the direct carbon emissions per capita of residents in 30 provinces, municipalities, and autonomous regions in China from 1999 to 2019. Among them, 19 kinds of energy consumption of residents’ life come from *China Energy Statistical Yearbook* in 2000-2019. The CO_2_ emission factor is derived from the *2006 IPCC Guidelines for National Greenhouse Gas Inventories*. The carbon emission coefficients of provinces and cities are derived from the *Emission Factors for China’s Regional Grid Baseline in 2019*. The total population, as well as urban and rural population comes from the *China Statistical Yearbook* in 2000-2019. It should be pointed out that due to the lack of relevant statistical data, the carbon emission data of the four regions of the Tibet Autonomous Region, Taiwan Province, Hong Kong, and Macao Special Administrative Region are not included in the investigated region. The relevant data of the Ningxia Hui Autonomous Region from 2000 to 2002 are missing, and the moving average method is used to supplement it. MSE is used for measuring the average error, which is relatively simple. It is used to measure the degree of change in the data. The smaller the MSE value, the higher the data prediction accuracy of the neural network model [19].

### 2.3 Carbon emissions prediction based on BP neural network model

BP neural network is a supervised learning algorithm, which consists of an input layer, hidden layer, and output layer. The full connection is formed between neurons of various layers [20]. According to the learning process, it is divided into a forward propagation and back propagation. The forward propagation is from the input layer to the hidden layer to the output layer. The back propagation is the signal propagating forward from the output layer. Each layer of transfer is limited by the weight value. The data information is processed by a combination of neuron activation function, hidden layer neuron quantity, and weight adjustment rules. Different network functions can be realized. Its specific structure is shown in Figure 1.

**Figure. 1.**
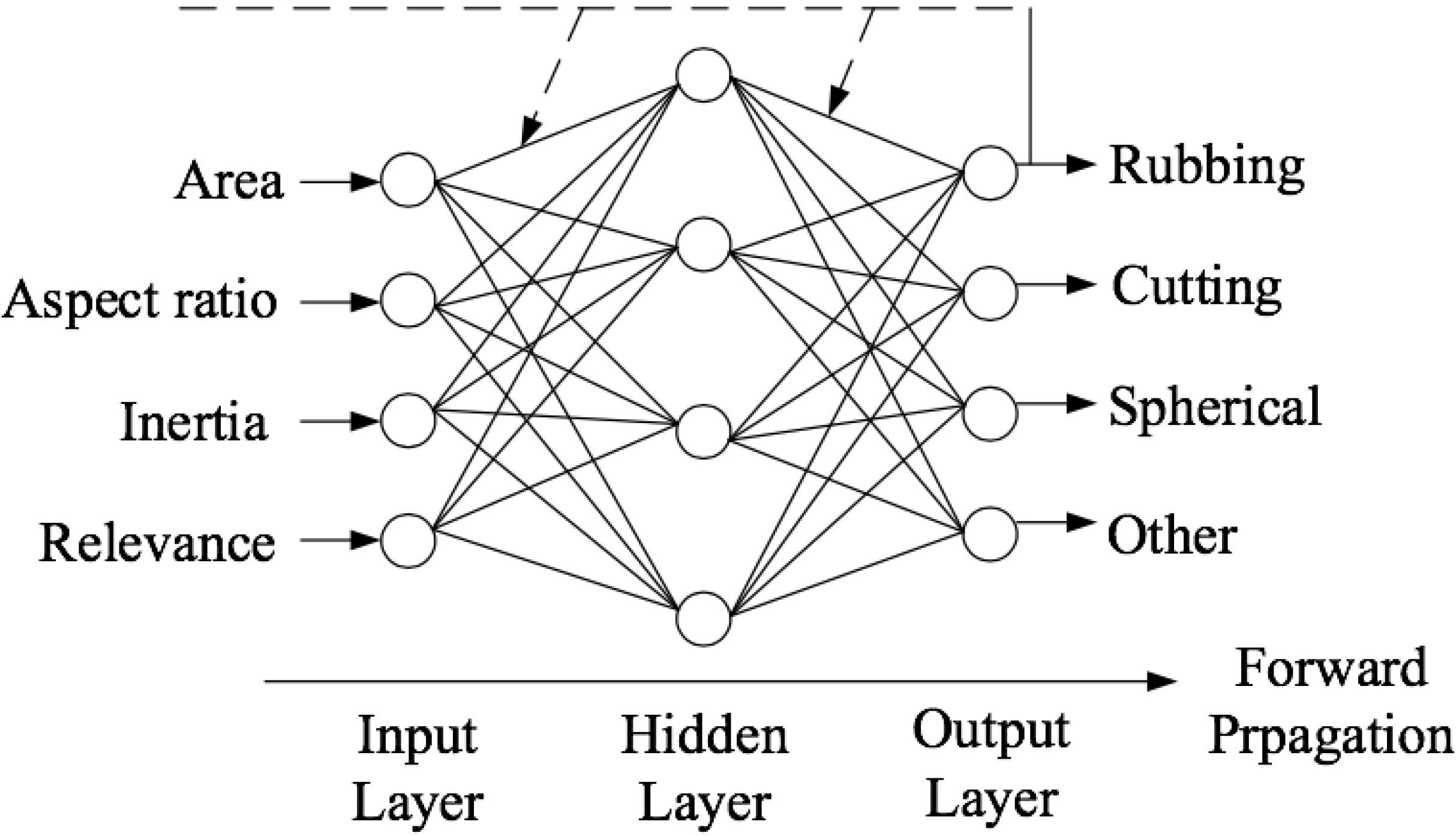
The schematic diagram of BP neural network structure

Based on the direct carbon emission data processing of Chinese provinces, the per capita carbon emission data of Fuzhou City is selected as the testing data of the neural network. The specific process is as follows: (1) Constructing training samples. All data samples are divided into 20 data samples. Since 1999, the last data every 5 years is the algorithm input value. The first 18 samples are the training set, and the last 2 are the testing set. (2) Carbon emission prediction and result output. For the already trained network, the Fuzhou carbon emission data in 2015-2019 are predicted and the results are output. Also, the residual error, relative error, and MSE are selected as the comparison standard of prediction accuracy.

### 2.4 Carbon emissions prediction based on RBF neural network model

RBF (Radial Basis Function) neural network is a three-layer feedforward neural network model with only one hidden layer. The input layer to the hidden layer uses a non-weighted connection, which can directly transfer the data to the hidden layer neural unit [21]. Figure 2 shows the structure of the RBF neural network model. It uses radial basis functions as the basis functions. The hidden layer is used to map the input data to the hidden layer space to complete the nonlinear transformation of the data. The output layer is often linearly transformed.

**Figure. 2.**
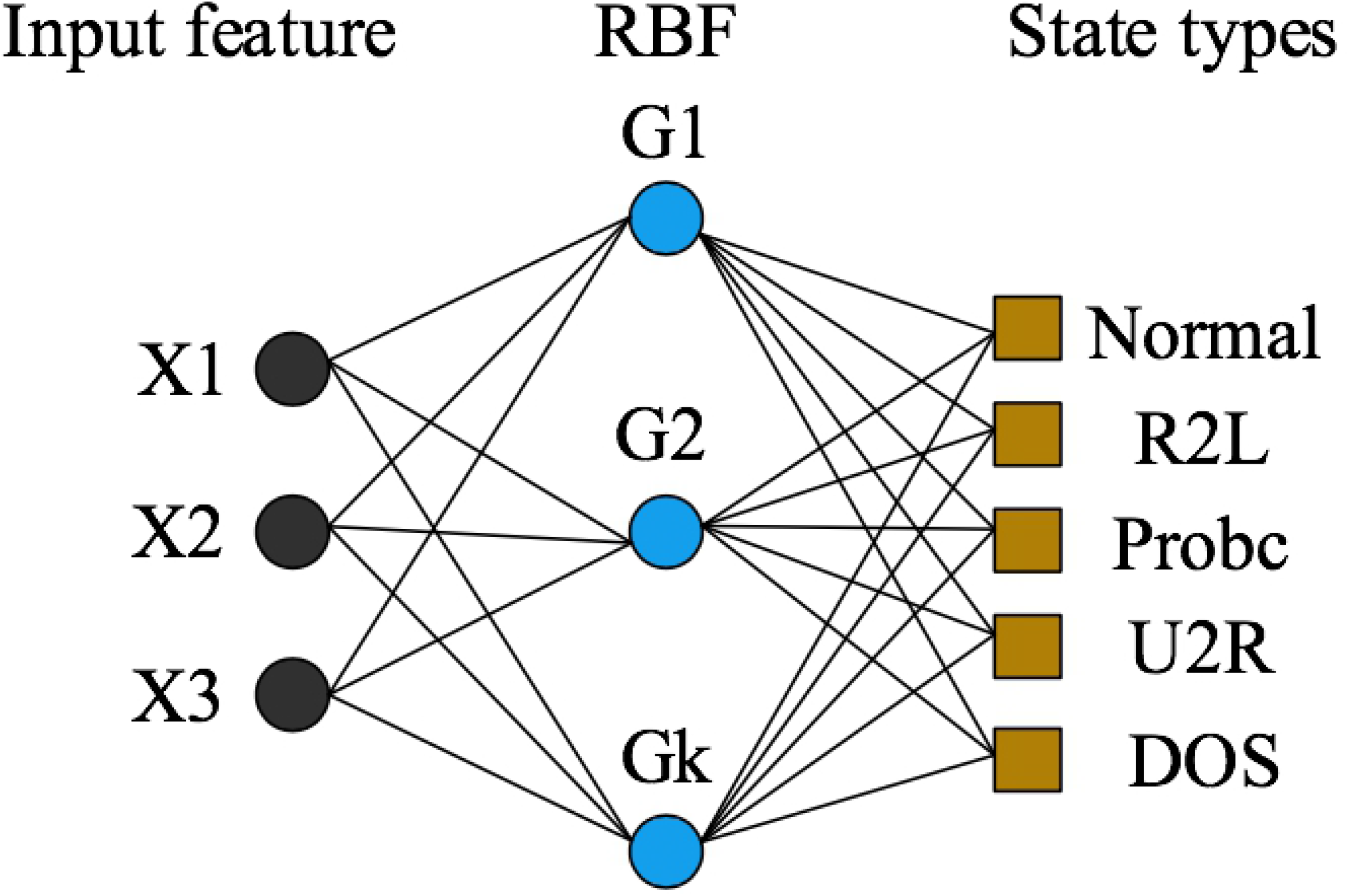
The schematic diagram of the RBF neural network structure

Based on the direct carbon emission data processing of Chinese provinces, the per capita carbon emission data of Fuzhou City is selected as the prediction purpose of the neural network. The specific process is as follows: (1) Sample data structure: The data is based on a 20×31 matrix. The data of other provinces except Fuzhou is used as the learning sample. Data from Beijing is used for testing. (2) The newrb function is used to set its network parameters such as allowable error, diffusion factor, and the number of neurons. On this basis, the same evaluation indicators as the BP neural network are adopted to output the results.

### 2.5 Carbon emission prediction based on Elman neural network model

Elman neural network is a dynamic feedback neural network, which is a neural network model with local memory and feedback capabilities. For the model, the convergence speed is good and the prediction accuracy is high. Therefore, it is adopted in many fields [22]. Compared with the above two network models, it has an additional undertaking layer with memory and feedback functions. The specific structure is shown in Figure 3. The undertaking layer can feed back to the hidden layer through data and make the network have the function of dynamic memory and feedback.

**Figure. 3.**
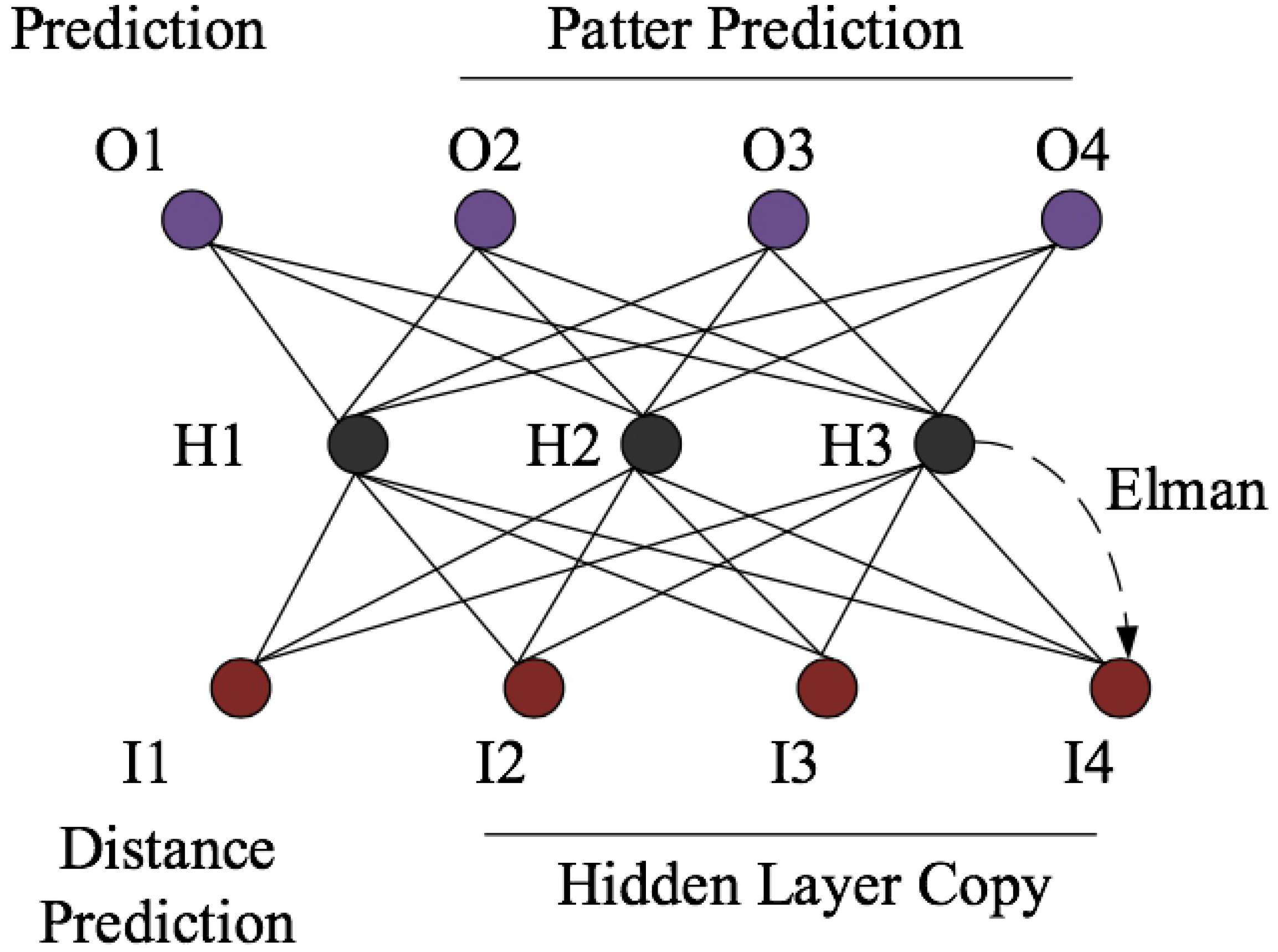
The schematic diagram of Elman neural network structure

Based on the direct carbon emission data processing of Chinese provinces, the per capita carbon emission data of Fuzhou City is selected as the prediction purpose of the neural network. The specific process is as follows: (1) Training sample determination: Consistent with the training sample method of BP neural network, the rolling prediction is realized in time series. (2) The elmannet function in the Matlab neural network toolbox is used to establish the Elman neural network. Also, the delay layer, hidden layer neuron size, training function, and other network parameters are set. (3) It is consistent with the time selection and measurement indicator of the BP neural network, and the results are output.

### 2.6 Carbon emission prediction based on GRNN neural network model

GRNN (Generalized Regression Neural Network) is a neural network algorithm under radial basis functions. There is strong curve mapping ability, flexible network structure, high fault tolerance, and fast learning speed in the algorithm. It has been applied in the construction of multiple network models [23]. The neural network can get a better prediction effect under the premise of a few samples. The specific structure is shown in Figure 4. In this investigation, 90% of the Fuzhou data in the residential carbon emission data is used as the training sample, and the rest is the testing sample. The measurement indicators and result output are similar to the BP neural network.

**Figure. 4.**
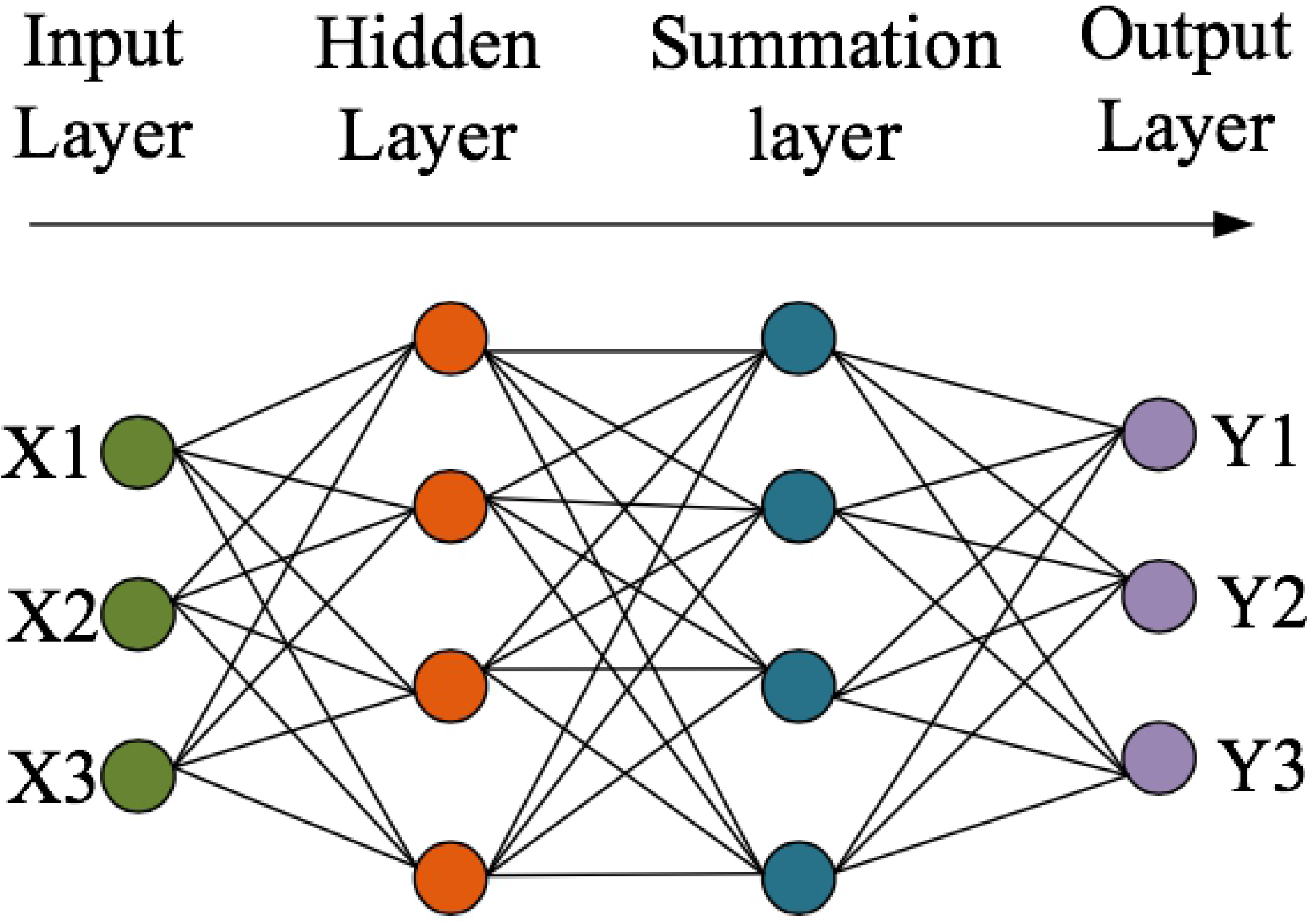
The schematic diagram of the GRNN neural network

### 2.7 System requirements and implementation

(1) System requirements: In terms of the basis, the system should have the basic functions of geographic information system software to implement basic map operations. In terms of parameters, it is necessary to call the Matlab software, modify the relevant parameters of the Elman neural network, set the predicted province and year, as well as display and save the carbon emission prediction results. In terms of display, thematic map creation functions are added to the system, including pie charts, bar charts, and graded coloring charts, more intuitively understanding the carbon emissions of various years and provinces as well as the composition of various energy consumption types. Also, it has other necessary functions for the production of related maps.

(2) System implementation: This system is developed under the C#.NET environment, based on ArcGIS Engine 10.2 combined with the neural network toolbox in Matlab R2014a. The C# language is derived from C and C++. While inheriting the powerful programming functions of C and C++ languages, the complex features of the two are eliminated. ArcGIS Engine is a complete set of embedded GIS component libraries and tool libraries that package ArcObjects. It is a complete class used by developers to build custom applications. The neural network toolbox in Matlab provides a practical technical method for the prediction of China’s provincial carbon emissions. Under the above development environment, this system calls the Elman neural network prediction model realized by using Matlab neural network toolbox in the form of the dll format, thereby realizing the integration of system functions.

## 3. Results and discussion

### 3.1 Total carbon emissions of residents in different years in different provinces

By calculating the data of different provinces and different years, the total carbon emission of residents shown in Figure 5 is obtained. From the figure, the total carbon emissions of residents in the seven provinces of Beijing, Tianjin, Hebei, Shanxi, Inner Mongolia, Liaoning, and Jilin in the northern region have not increased much from 2000 to 2010, but the growth rate has risen linearly after 2010. As far as Tianjin is concerned, compared with 2010, the total carbon emissions of residents in 2018 increases by 4 times. In the southern regions of Jiangsu, Zhejiang, Anhui, Fujian, and Jiangxi, the total carbon emissions have increased, but the growth rate has slowed distinctly. Among them, the fluctuations in Shanghai and Shandong provinces are relatively large, with a clear trend of increasing first and then decreasing. It is closely related to local environmental protection policies. For the provinces of Shaanxi, Gansu, Ningxia, Qingdao, and Guizhou in the northwestern region, there is a trend of a decrease after a short rise and then a slow rise.

**Figure. 5.**
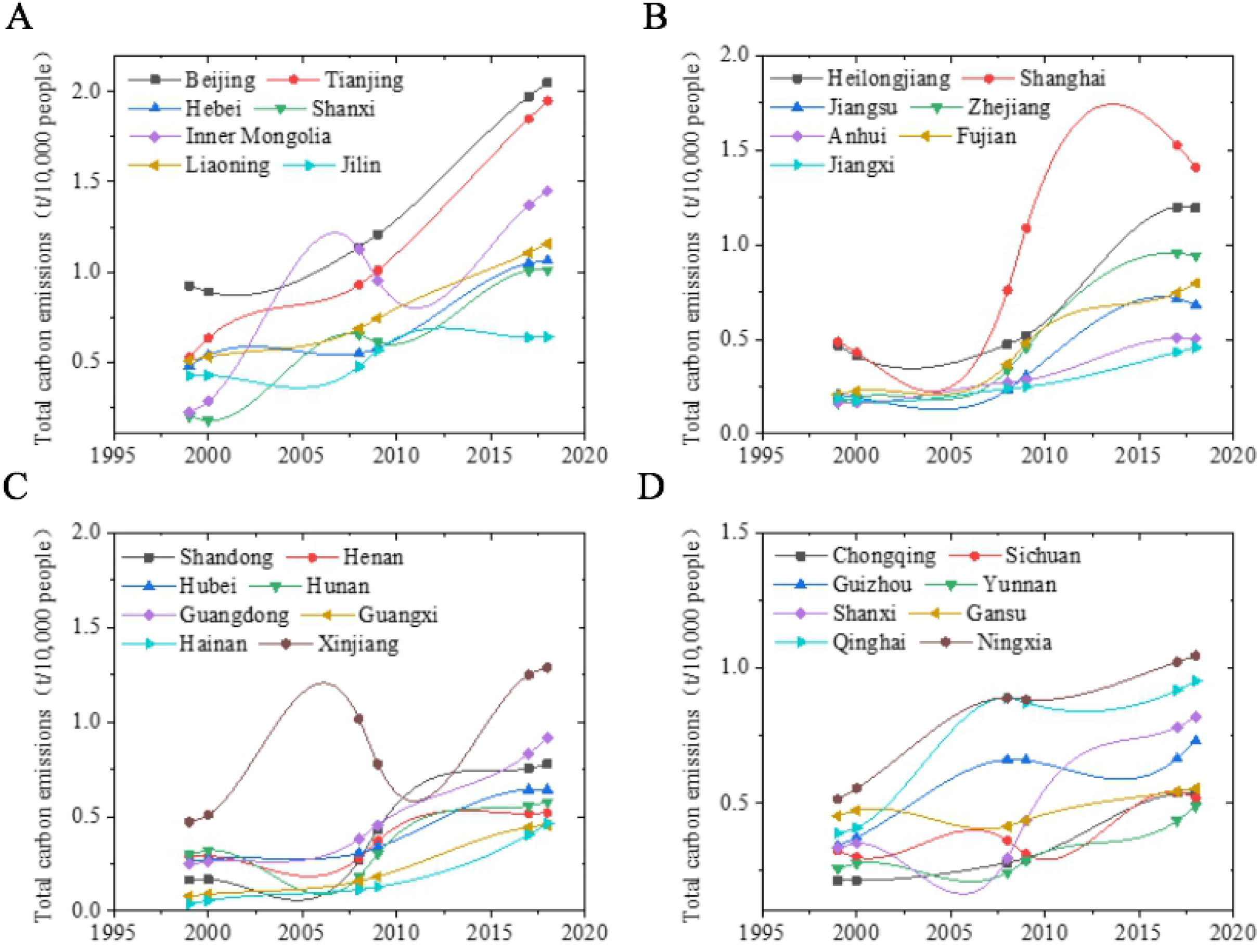
Total carbon emissions in different years in different provinces

### 3.2 Prediction results of different carbon emission prediction models in different provinces

Different carbon emission prediction models are used to predict the data from 2015 to 2019 in the four core geographic regions of China, Beijing, Zhejiang, Guangdong, and Shaanxi. The results are shown in Figure 6. From the figure, except for the large error between the predicted values of 2018 and 2019 obtained by using the GRNN neural network, the changing trends of the predicted carbon emissions of other neural networks are almost consistent with the actual carbon emissions. The prediction results are relatively ideal. Compared to the BP neural network model, the RBF neural network model has a simple structure without requiring repeated training. But the results show that the neural network has low prediction accuracy and large errors. Compared to the BP and RBF neural network models, the Elman neural network has higher accuracy in predicting carbon emission data in four regions and has a better prediction effect.

**Figure. 6.**
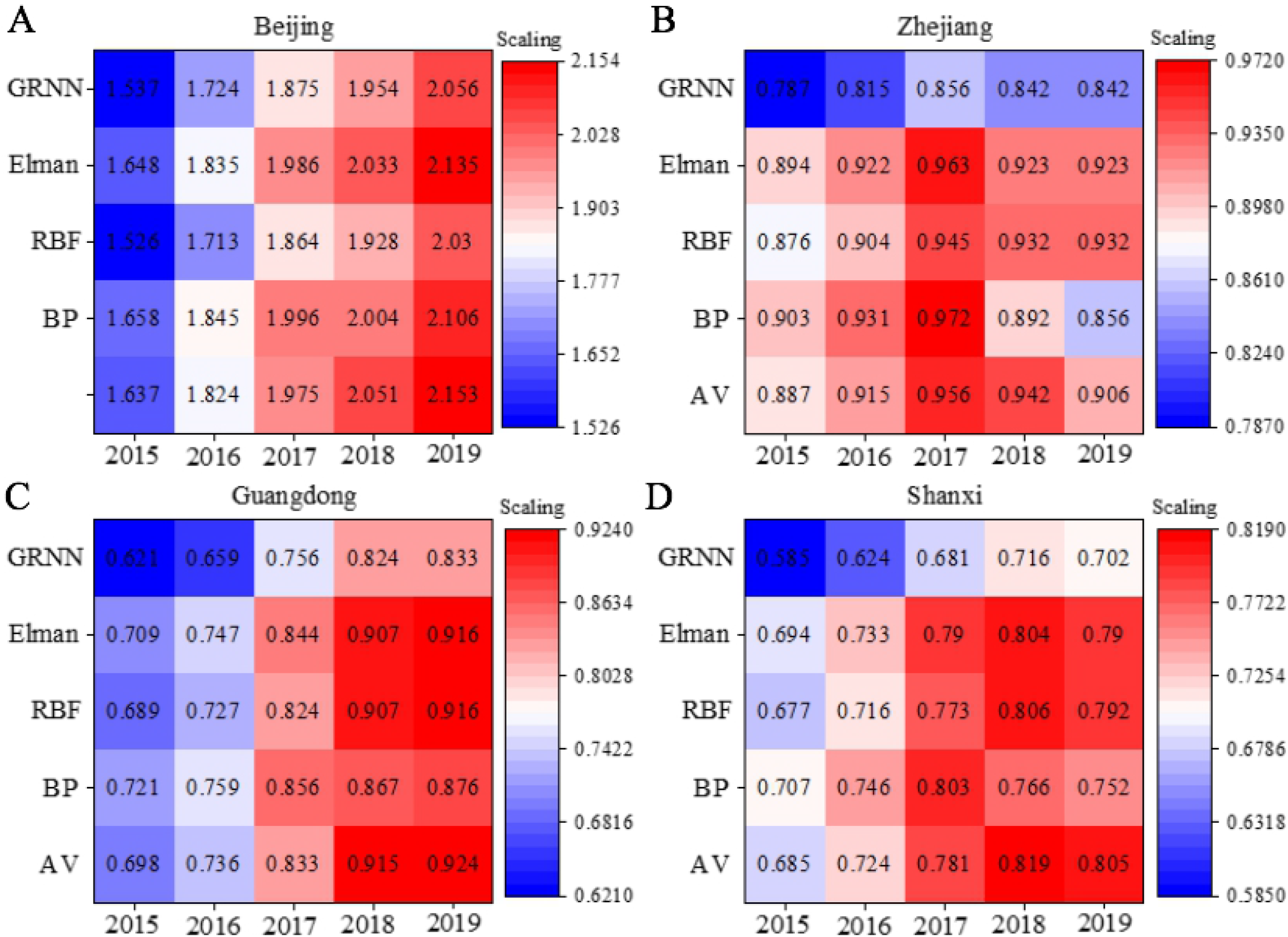
Prediction results of different carbon emission prediction models in different provinces

### 3.3 Performance evaluation of different carbon emission prediction models

The above data are analyzed for errors and relative errors. Table 1 shows the results. From the table, compared to the RBF neural network, BP neural network requires multiple trainings according to experience to determine the better network in different years. Therefore, a large prediction error occurs during the training process. Compared to the Elman neural network, the RBF neural network model has insufficient robustness and greater randomness. Therefore, the prediction accuracy is slightly inferior to the Elman neural network. Based on the above results, the effect of the Elman prediction model is optimal.

**Table. 1.**
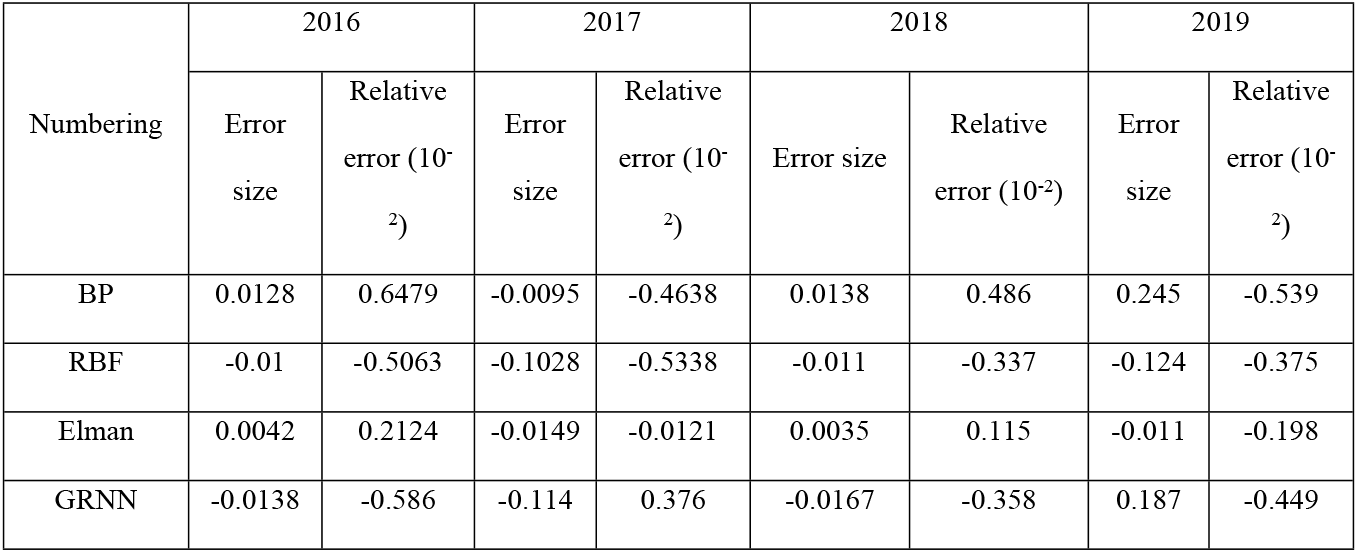
Performance evaluation of different carbon emission prediction models

Figure 7 shows the prediction performance evaluation of different carbon emission models. From the figure, in terms of MSE, the smaller the value, the higher the accuracy of the model. The MSE of the BP neural network model after deep learning is 0.0252. Compared with other models, its accuracy performance is the worst. The optimal is the Elman carbon content prediction model. Compared to the BP neural network model, accuracy is improved by 55.93%. In terms of average MSE, the larger the value, the better the model prediction performance, which is consistent with the accuracy results. Compared to the BP neural network model, the prediction performance of the Elman network model is improved by 19.48%. In terms of the goodness of fit, the Elman network model has not improved much compared to other network models.

**Figure. 7.**
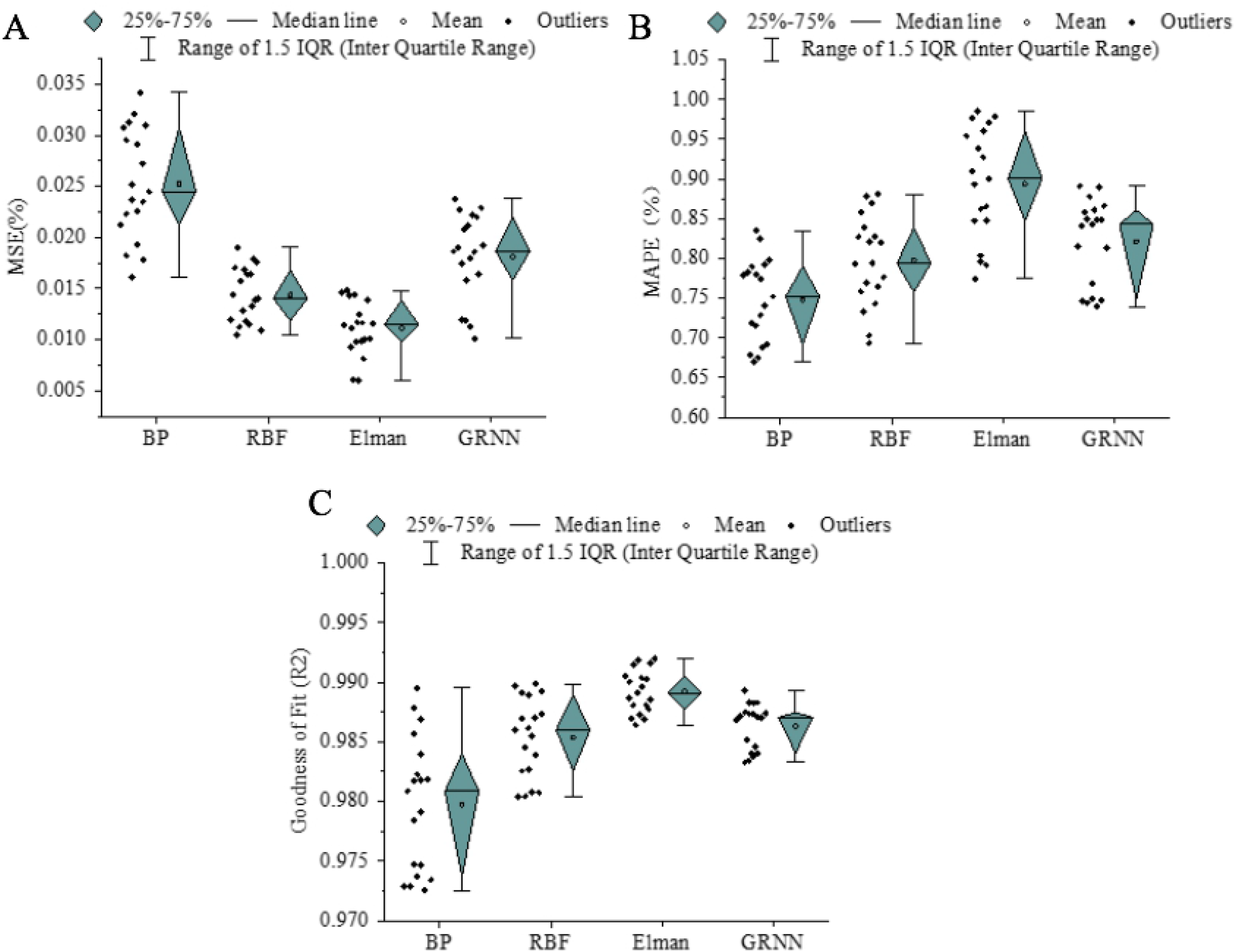
Prediction performance evaluation of different carbon emission models

### 3.4 Prediction results of direct carbon emissions from Chinese residents in the future

Based on the above model comparison, the Elman carbon emission prediction model is selected to predict the direct carbon emissions of Chinese residents from 2020 to 2035. The results are shown in Table 2. From the table, without considering other influencing factors (especially policies), the direct carbon emissions of Chinese residents in the next 15 years will still show a steady growth trend in the next 5 years. After 2027, direct carbon emissions per capita will show a slight downward trend. Also, in 2032, the carbon emissions per capita will be 0.8882t/10,000 people. After reaching a weak peak, it will show a slight decrease. Based on the above results, it can be concluded that in theory it is expected to reach the peak of carbon emissions around 2027 to 2032.

**Table. 2.**
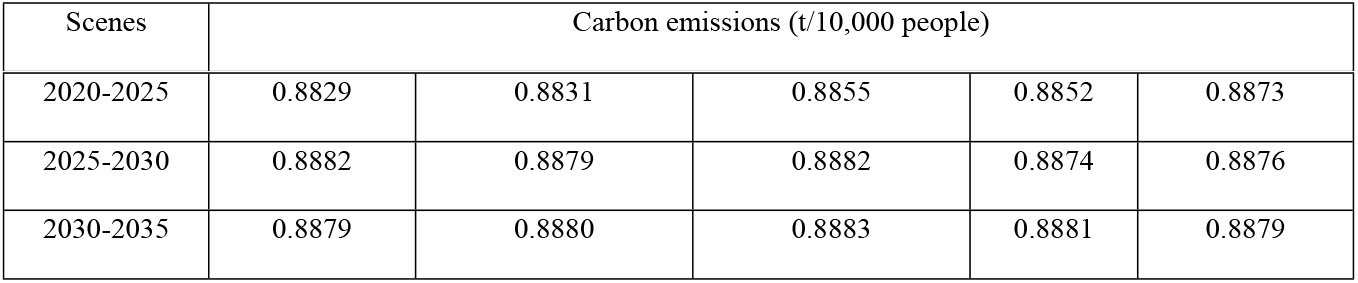
Prediction results of Chinese residents’ direct carbon emissions from 2020 to 2035

| Scenes | Carbon emissions (t/10,000 people) |
| --- | --- |

|  |  |  |  |  |  |
| --- | --- | --- | --- | --- | --- |
| 2020-2025 | 0.8829 | 0.8831 | 0.8855 | 0.8852 | 0.8873 |
| 2025-2030 | 0.8882 | 0.8879 | 0.8882 | 0.8874 | 0.8876 |
| 2030-2035 | 0.8879 | 0.8880 | 0.8883 | 0.8881 | 0.8879 |

## 4. Conclusion

Under the severe pressure of energy saving and emission reduction, through the national statistical data information, based on the artificial neural network model, mathematical statistics and deep learning methods are used to learn and analyze the carbon emission data of various provinces in China from 1999 to 2019. Different neural network models are selected, and the carbon emission data of Fuzhou are used as testing data to make predictions. The feasibility of the neural network model to predict carbon emission data is verified. Also, the prediction performance is compared to select a better network, embedding it into the direct carbon emission prediction system of Chinese provincial residents. It provides an effective operating platform for carbon emission analysis and prediction. Compared with other models, Elman neural network has higher accuracy and smaller error in carbon emission prediction. Compared to the BP neural network, the accuracy is improved by 50%, and the prediction performance is improved by nearly 20%. It is better to use this prediction model to predict residents’ carbon emissions. Although the advantages and disadvantages of different models are analyzed as much as possible, there are still many disadvantages in some places due to limited time and energy: (1) Affected by factors such as the availability of statistical data over the years and different statistical calibers, the calculated direct carbon emission data of residents in various years and provinces will inevitably produce errors. It leads to problems with the conclusions reached. (2) At present, more empirical methods are used to obtain a more ideal network model. The network testing process is relatively cumbersome and the workload is large. In this process, subjective factors will have a certain impact on network performance. Next, in-depth investigations will be conducted in these two aspects, quickly and efficiently predicting and analyzing the carbon emissions of residents.

